# Regulation of traction force through the direct binding of Basigin (CD147) and Calpain 4

**DOI:** 10.1101/2023.03.06.531406

**Authors:** Bingqing Hao, Jacob DeTone, Mia Stewart, Savannah Kozole, Karen A. Beningo

## Abstract

Traction force and mechanosensing (the ability to sense the mechanical attributes of the environment) are two key factors that enable a cell to modify its behavior during migration. Previously, it was determined that the calpain small subunit, calpain 4 (CapnS1), regulates the production of traction force independent of its proteolytic holoenzyme. A proteolytic enzyme is formed by calpain 4 binding to either of its catalytic partners, calpain 1 and 2. To further understand how calpain 4 regulates traction force, we used two-hybrid analysis to identify more components of the traction pathway. We discovered that basigin, an integral membrane protein and a documented inducer of matrix-metalloprotease (MMP), binds to calpain 4 in two-hybrid and pull-down assays. Traction force was deficient when basigin was silenced in MEF cells, and this deficiency was also reflected in the defect in substrate adhesion strength. Unlike Capn4^-/-^ MEF cells, the cells deficient in basigin had normal mechanosensing abilities. Together, these results implicate basigin in the pathway in which calpain 4 regulates traction force independent of the catalytic large subunits.

## INTRODUCTION

Cell migration is essential for various normal and abnormal physiological processes, including embryonic development, wound healing, immune responses, and cancer metastasis. In addition, cell migration is significant to technological applications such as tissue engineering [1–4]. Although numerous studies have been conducted to extend our understanding of how the complex process of cell migration is regulated, the mechanism remains to be fully defined.

Focal adhesions function dynamically in cell migration, specifically in biophysical terms of transmitting both traction forces and mechanosensing between the actin cytoskeleton and extracellular matrix (ECM) [5–8]. Calpains have been implicated in the study of cell migration since calpain proteases localize to focal adhesions and play essential roles in the turnover of several focal adhesion components [9–12]. The two best characterized calpains, μ-calpain and m-calpain, both contain a distinct 80 kDa catalytic large subunit (calpain 1/CAPN1 and calpain 2/CAPN2, encoded by *Capn1 and Capn2* genes, respectively) and a common 28kDa small subunit (calpain 4/CAPNS1/CAPN4, encoded by *Capn4* gene) [12]. Inhibiting calpains through the overexpression of the endogenous inhibitor calpastatin and the use of pharmacological inhibitors leads to the inhibition of both adhesive complex disassembly and actinin localization to focal contacts when cells are attached to glass coverslips [10].

Calpains are known to be regulated post-translationally through phosphorylation events, the endogenous inhibitor, and interactions with the regulatory small subunit. The protein phosphatase 2A (PP2A) is identified as a calpain phosphatase of μ-calpain and m-calpain and can directly dephosphorylate both heavy chains. The dephosphorylation by PP2A inactivates μ-calpain and m-calpain, resulting in the suppression of migration in lung cancer cells [13] (Xu and Deng, 2006). The small subunit was historically considered to mainly serve a regulatory function for calpain holoenzymes [12]. However, a finding that *Capn4^−/–^* embryonic fibroblasts present abnormal organization of focal adhesions, reduced rates of cell migration, and delayed retraction of membrane projections implicates the small subunit in the regulation of cell migration [11]. In addition, we have found that traction force was attenuated by the knockout or knockdown of the calpain small subunit but not by the large subunits. In contrast, all subunits are required for mechanosensing. This study implicated only the small subunit functioning as an independent entity in regulating traction force [14, 15].

To gain insight into how the calpain small subunit regulates the production of traction force, we screened for its direct binding partners. In this study, we used the entire Capn4 gene as bait in a yeast two-hybrid assay. From a screen of the whole mouse embryonic genome, we identified the protein basigin as a direct binding partner of calpain 4.

Basigin (*Bsg*), also known as CD147 or EMMPRIN, is a heavily glycosylated transmembrane protein belonging to the immunoglobulin (Ig) superfamily [16, 17]. Basigin has been found to play roles in various biological processes and in the progression of cancer. Mice deficient in the basigin gene have abnormal embryogenesis, spermatogenesis, and fertilization [18–20]. Knock-out mice of the *Bsg* gene display abnormalities in vision and insensitivity to irritating odor [21, 22]. Basigin is commonly overexpressed in many tumors [23–25] and is implicated in almost all types of cancer [24]. On the surface of tumor cells, basigin was found to stimulate the production of matrix metalloproteinases (MMPs) in adjacent fibroblasts, resulting in enhanced tumor invasion, which led to the name EMMPRIN [26, 27].

Basigin is known to interact (directly or indirectly) with several proteins, including MCT1, MCT2, integrin-β1, cyclophilin, caveolin-1, annexin-2, and ubiquitin C [24, 28–30]. Many of its known interacting proteins are implicated in cell migration and secretion. Basigin’s functions in tumorigenesis and cell migration render it a reasonable target candidate for elucidating how calpain 4 regulates the production of traction force.

In this study, basigin was identified as a binding partner for calpain 4 using the yeast two-hybrid assay and subsequently verified through immunoprecipitation. Furthermore, we found that upon basigin knockdown, traction force was significantly reduced, and these cells were defective in substrate adhesion. Surprisingly, MEFs in which basigin is silenced can respond to localized stimuli, sense the stiffness of substrates like wild-type cells, suggesting basigin functions only in the production of traction stress with calpain 4 but not in mechanosensing, where calpain 4 is needed. These results implicate basigin in the same pathway that calpain 4 functions to regulate the production of traction force, a path that is independent of the catalytic activity of the holoenzyme.

## EXPERIMENTAL PROCEDURES

### Cell Culture

Mouse embryonic fibroblasts expressing a defective small calpain subunit have been described previously [11, 31](Arthur et al., 2000; Dourdin et al., 2001), and are referred to as *Capn4^−/–^* cells in this study. Mouse embryonic fibroblasts (MEFs), Capn4–/– cells, and 293T cells were used in this study. MEFs were purchased from ATCC. 293T cells were a gift from Dr. Xiangdong Zhang, University of Buffalo. MEFs and *Capn4^−/–^* cells were maintained in Dulbecco’s Modified Eagle’s Medium-high glucose (Sigma) supplemented with 10% fetal bovine serum (FBS) (Hyclone) and 1% penicillin/streptomycin/glutamine (PSG, Gibco) and incubated at 37°C under 5% CO2 in a humidified cell culture incubator. 293T cells were maintained and passed similarly, except that the media contained 1% PS instead of PSG. Cells were passaged by trypsinization using 0.1% trypsin-EDTA (0.25% trypsin-EDTA diluted in HBSS, Gibco) and then diluted and transferred into new culture dishes. Passages were limited to eight per cell line.

### Cloning of CAPN4 and Yeast Two-Hybrid Assay

Full-length *Capn4* was amplified by PCR from a pEGFP-capn4 plasmid under the following conditions: 30 cycles of 98 °C for 10 sec followed by 68 °C for 1 min using PrimeSTAR HS DNA Polymerase with GC buffer (Takara, R044A). The primers used to insert *Capn4* into pCWX200 and pLexA were the forward primer, 5’-ATCGGGATCCTTATGTTCTTGGTGAATTCGTTCTTGAAGG-3’, and the reverse primer, 5 5’-ACCGCTCGAGTCAGGAATACATAGTCAGCTGCAGCCAC-3’. PCR products were resolved on 1% agarose gels and visualized by ethidium bromide (1% solution, Fisher) staining. The resolved bands were then purified using a Qiaquick gel extraction kit (Qiagen, 28706). Purified PCR products, pCWX200 and pLexA were incubated with XhoI and BamHI (New England Biolabs) at 37°C for 4 hours in 1X buffer 3 supplemented with 1% BSA. The double-digested PCR products and plasmids were again purified with the Qiaquick gel extraction kit. To insert *Capn4* into pCWX200 and pLexA, the double-digested Capn4 PCR product was ligated with the double-digested pCWX200 or pLexA using the LigaFast Rapid DNA Ligation System (Promega, M8226). Successful insertions were confirmed by DNA sequencing (Applied Genomics Technology Center, Wayne State University). Bait plasmids were provided to ProteinLinks Inc. (Pasadena, CA) for the yeast two-hybrid assay. The candidates obtained via two-hybrid were identified by DNA sequencing (Applied Genomics Technology Center, Wayne State University).

### Small Interfering RNA (siRNA) Nucleofection

Wild-type MEFs were used for the selective silencing of *Bsg* via siRNA. The knockdown was generated through transient transfection with either control siRNA oligonucleotides or siRNA oligonucleotides targeting the *Bsg* gene using the siGENOME SMARTpool system (Dhamacon). The siRNA oligonucleotides targeting the *Bsg* gene were: GAUUGGUUCUGGUUUAAGA, CAUCAGCAACCUUGACGUA,GCAAGUCCGAUGCAUCCUA,GGACAAGAAUGUACGC CAG. Nucleofection was performed using the Amaxa MEF2 Nucleofector Kit (Lonza) following the manufacturer’s suggested protocol. Briefly, MEF cells were trypsinized with 0.1% Trypsin-EDTA and collected by centrifuging at 2000 rpm for 5 min. Collected cells were then resuspended in an appropriate volume of the mixture, which included the MEF2 nucleofector solution, up to 5 μg of control siRNA or siRNA targeting the Bsg gene, and supplement 1 provided with the kit. The total volume of the MEF2 Nucleofector solution, supplement 1 mixture, and siRNA was 100 μl and was transferred to an electroporation cuvette. The cuvette was then inserted into the Nucleofector II system (Amaxa) and the program MEF A-023 was run. 500 μL of RPMI-1640 medium (Sigma) was immediately added to the cuvette after the program was run to prevent cell damage. Nucleofected cells were then seeded according to the requirements of the following procedures. Inhibition of basigin expression reaches a maximum at 36 hrs after nucleofection.

### Protein Extraction and Western Blotting

Proteins were extracted from each cell line using triple-detergent lysis buffer (TDLB): pH 8, 50 mM Tris-HCl, 150 mM NaCl, 1% NP-40, 0. 5% sodium deoxycholate, 0. 1% SDS, with added Protease Inhibitor Cocktail (Sigma) and HaltTM Phosphatase Inhibitor Cocktail (ThermoFisher). An 80% confluent 100 mm culture dish (NuncTM) was placed on ice and washed twice with ice-cold phosphate-buffered saline (PBS), then incubated for 25 minutes with 300 μl TDLB on ice. Lysed cells were collected into 1. 5 ml tubes using an ice-cold cell lifter and centrifuged at 13, 000 rpm for 10 minutes to remove cell debris. Protein concentration was determined using the Bio-Rad DC protein assay kit, following the manufacturer’ s instructions. Proteins were isolated from cell lines of MEFs, capn 4-/-cells, MEFs transfected with control siRNA, and MEFs transfected with siRNA targeting the Bsg gene. Twenty micrograms of protein were loaded onto a 4-20% gradient Tris–HEPES–SDS precast polyacrylamide gel system (Pierce) and resolved at 100 V for 1 hour. Proteins were then transferred onto an Immuno-Blot PVDF membrane (Bio-Rad) using a Trans-blot Turbo Transfer System (Bio-Rad) at 25 V for 15 minutes. After transfer, the membrane was blocked for 1 hour with 5% milk in 1 X phosphate-buffered saline (PBS) – 0. 1% Tween (0. 1% PBS/T) for the basigin p-antibody, or in 5% milk in 1 X Tris-buffered saline (TBS)-0. 1% Tween (0. 1% TBS/T) for anti-actinin, anti-FLAG, and anti-HA antibodies. The membrane was then incubated overnight at 4 ° C with primary antibodies with gentle agitation. The primary polyclonal or monoclonal anti-basigin antibody (sc-9757, B-5, Santa Cruz) was diluted at 1: 800 in 5% milk in 0. 1% PBS/T. Anti-α-actinin antibody (A 5044, Sigma) and monoclonal anti-FLAG antibody (F 1804, Sigma) were diluted at 1: 500 in 5% milk in 0. 1% TBS/T. Monoclonal polyclonal anti-HA antibody (MMS-101 P, Covance) was diluted at 1: 1000 in 5% milk in 0. 1% TBS/T. After washing three times for 20 minutes each with 0. 1% PBS/T, the membrane was incubated for 1 hour at room temperature with secondary antibodies. HRP-linked anti-goat IgG (sc-2020, Santa Cruz) at 1: 2000 dilution was used for the p-anti basigin antibody, while HRP-linked anti-mouse antibody (ThermoFisher) at 1: 10, 0.1 dilution was used for anti-actinin, anti-FLAG, and anti-HA antibodies. For some blots, total protein was detected using No-Stain total protein labeling reagent (Invitrogen). After washing three times for 20 minutes each, detection was performed using ECL Plus Western Blotting Detection Reagents (Amersham), and visualization was done with an iBright 1500 (ThermoFisher).

### Cloning of CAPN4 and BSG, and Immunoprecipitation

Full-length *Capn4* was amplified by PCR from the pEGFP-capn4 plasmid and inserted into a pFLAG-CMV vector. The BSG gene, lacking the sequence for the N-terminal 100 amino acids, was amplified from the pJG4-5-BSG vector, resulting from the yeast two-hybrid assay, and inserted into a pCDNA3 vector together with an HA sequence. The primers used for amplification of *Capn4* were: forward primer 5’-CCCAAGCTTATGTTCTTGGTGAATTCG-3’ and reverse primer 5’-CCGGGATCCTCAGGAATACATAGTCAGCTGC-3’. Basigin was amplified from a plasmid generously provided by Dr. Judith Ochrietor, University of North Florida. The primers used for amplification of *Bsg* were forward primer 5’-CGCGGATCCATGGAAGGGCCACCCAGGATCAA-3’ and reverse primer 5’-CCGCTCGAGTCAGGTGGCGTTCCTCTGG-3’. Successful insertions were confirmed by sequencing (Applied Genomics Technology Center, Wayne State University).

293T cells were co-transfected with 10μg of Flag-tagged Capn4 vector (full length) and 10μg of HA-tagged Basigin containing vector. 20 hours after the transfection, cells were harvested, and the immunoprecipitation assay was performed. Cells were lysed with ice-cold 1X lysis buffer (50 mM HEPES-NaOH, pH 7.5, 100 mM NaCl, 0.5% NP-40, 2.5 mM EDTA, 10% glycerol, 1 mM DTT) with Protease Inhibitor Cocktail (Sigma) and HaltTM Phosphatase Inhibitor Cocktail (Thermo) and then collected and pelleted by centrifugation. To 500 μg of cell lysate, 10 μg of anti-FLAG antibody or anti-HA antibody was added, and then the lysates were incubated for 1 hour at 4°C. 20 μl of Protein A/G PLUS Agarose (sc-2003, Santa Cruz) was added and incubated at 4°C on a rocker platform overnight. The immunoprecipitate was collected by centrifugation, and the pellet was washed four times with 1X lysis buffer. After the final wash, the pellet was resuspended in 40 μL of electrophoresis sample buffer. The sample was boiled for 3 minutes and analyzed by SDS-PAGE with corresponding antibodies.

### Digestion of Glycosyl Units from Basigin

Cell lysates were incubated with peptide-N-glycosidase (PNGase) (New England Biolabs) according to the manufacturer’s instructions. This involved incubating 9 μL of lysate with 1 μL of 10X Glycoprotein Denaturing Buffer at 95 °C for 10 minutes, followed by 5 minutes on ice. Samples then received 2uL 10X Reaction buffer, 2uL 10% NP-40, 6uL dH2O and 1uL PNGase (final conc of enzyme at 20U/uL) and incubated at 37°C for 2hr. Loading dye was added, and the samples were heated to 95°C to stop the reaction. Samples were run on SDS-PAGE and blotted as above.

### Preparation of Polyacrylamide Substrates

A series of polyacrylamide substrates of different stiffnesses was prepared as described previously [32]. Briefly, a flexible 75 μm × 22 mm polyacrylamide substrate was prepared in a cell culture chamber dish and embedded with 0.2 μm fluorescent microbeads. The acrylamide (acryl, Bio-rad) concentration was fixed at 5% while N, N-methylene-bis-acrylamide (bis, Bio-rad) varied from 0.04% to 0.1% to attain different stiffnesses of the substrates. The substrates were then coated with 5 μg/cm² fibronectin (Sigma) at 4°C overnight by cross-linking with Sulfo-SANPAH (Thermo). Cells were seeded onto the substrates overnight before TFM or mechanosensing assays. The 5%/0.08% Acryl/Bis substrates (*E*=1.41 kPa) were used in traction force microscopy (TFM), the 5%/0.1% Acryl/Bis substrates were used in the localized mechanosensing assay, and 5%/0.1% Acryl/Bis (hard) (*E*=2.11 kPa) and 5%/0.04% Acryl/Bis (soft) (*E*=0.41 kPa) substrates were used for the cell adhesion assay.

### Traction Force Microscopy

Flexible polyacrylamide substrates of 5%/0.08% Acryl/Bis coated with 5 μg/cm^2^ fibronectin were prepared as described above. Cells were seeded onto the flexible polyacrylamide substrates. Overnight cultures were imaged as described previously [32]. Briefly, for a single cell, three images were taken under a 40X objective lens: a bright-field image of the cell, an image of the fluorescent beads with the cell on the substrate, and another image of the fluorescent beads after a microneedle had removed the cell. DIM (program designed by Dr. Yu-li Wang) was used to calculate bead displacement with or without the cell and the cell and nuclear boundaries. These data were used to generate and render traction stress values by using an algorithm courtesy of Dr. Micah Dembo (Boston University) as described previously [33]. Images of 14-22 cells for each cell line were collected.

### Mechanosensing Assays

Flexible polyacrylamide substrates of 5/0.1% Acryl/Bis coated with fibronectin were prepared as described above. Cells were seeded onto the substrates and allowed to adhere overnight under regular cell culture conditions. As described previously [15] a cell was monitored for 10 minutes to track its migration trajectory before a blunted microneedle was pressed onto the substrate in the direction the cell was migrating, thereby generating a pushing force on the cell. The pushing force would release the tension on the substrate, creating a localized soft spot. Images were taken of cells using a 40X objective lens at 3-minute intervals for 1 hour to record the migrating trajectories of the cells. If a cell responds to the pushing force applied by the needle by changing trajectory or rounding up, this is a positive response. If a cell continues to migrate on the same trajectory, it does not sense the change in the microenvironment and is recorded as not responding. For each cell line, 6-10 cells were observed.

To explore the effect of global compliance, as opposed to the localized change in compliance tested above, on cellular morphology, polyacrylamide substrates of stiffness of 5%/0.1% Acryl/Bis (hard) and 5%/0.04% Acryl/Bis (soft) were made as described above. After solidification, the substrates were coated with 5 μg/cm^2^ fibronectin. Cells were coated onto the substrates and allowed to adhere overnight under regular cell culture conditions before the images were recorded with a 10X objective lens. The number of round cells was counted from zoomed images from six random fields for each cell line on both substrates with different stiffnesses. The cell numbers were plotted into box and whisker plots.

### Cell Adhesion Assay

A centrifugation assay was used to measure cell-substrate adhesiveness. This assay was performed using the method described by Guo et al. [34], with slight modifications. Briefly, a hole was drilled in an air-tight culture dish (Pall Corporation), and a coverslip was attached to the culture dish. 5%/0.08% Acryl/Bis substrates were prepared on coverslips as described above and then coated with 5 μg/cm² fibronectin. 2.5 × 10^4 cells were seeded onto fibronectin-coated substrates and allowed to adhere for 30 minutes at 37°C. After incubation, the chambers were inverted and centrifuged for 5 minutes at 1800 × g. Ten random fields of cells were counted for each cell line right away after centrifugation. Percentages of cells before and after centrifugation are expressed in column graphs.

### Immunofluorescence

After being flamed, No. 1.5 glass coverslips (Fisher) were attached to chamber dishes. Then, they were coated with 5 μg/cm² fibronectin (Sigma) at 4°C overnight and blocked with 1% BSA in PBS at 4^°^C overnight. Cells were seeded onto coverslips and allowed to attach overnight under incubation at 37°C with 5% CO2 in a humidified cell culture incubator. The cells were then fixed and permeabilized with the following steps: firstly, incubate for 10 min with 2.5% paraformaldehyde in 1X PBS at 37^°^C; then incubate with 2.5% paraformaldehyde in 1X PBS with 0.1% Triton X-100 at 37^°^C; followed by incubation for 5 min with 0.5 mg/ml NaBH_4_ solution. After fixation and permeabilization, cells were blocked with 5% BSA in PBS for 1 hour at room temperature. Then they were incubated with basigin antibody (sc-9757, Santa Cruz) at a 1:250 dilution for 3 hours at room temperature. Following three 15-minute washes, Alexa Fluor 546 anti-goat secondary antibody was added at a 1:500 dilution in 5% BSA for an incubation of 1 hour at room temperature. After the final washes (3 x 15 min each), mounting media (pH=7.8, 0.1% PPD, 1X PBS, 50% glycerol, 30% Q-H_2_O) was added. Images were taken with appropriate filters for GFP signals.

### Cell Migration Assay

After being flamed, No. 1.5 square glass coverslips (Fisher) were attached to chamber dishes and coated with 5 μg/cm² fibronectin (Sigma) diluted in 50 mM HEPES at 4°C overnight. Cells were then seeded onto the coverslips and allowed to attach overnight under incubation at 37°C with 5% CO2 in a humidified cell culture incubator. Images were taken to describe the migration trajectory of a single cell for 2 hours at 2-minute intervals with a 40X objective lens. All the collected images for one cell were imported into the custom-built dynamic image analysis system software (DIM, Y-L. Wang) to calculate the linear speed and persistence of each cell. 15-18 cells were observed for each cell line.

### Microscopy

Images of all experiments described above were acquired using an Olympus IX81 ZDC inverted microscope equipped with a custom-built stage incubator to maintain cells at 37°C under 5% CO2 for live-cell imaging. A SPOT Boost EM-CCD-BT2000 back-thinned camera (Diagnostic Instruments Inc., Sterling Heights, MI) was also used. The camera was run by IPLab software (BD Biosciences).

### GEPIA2-Guided Pan-Cancer Analysis of CAPN4 and Bsg

Given the propensity of CAPN4 and Bsg overexpression in tumor progression, we explored their correlation in various tumors of interest. The GEPIA2 (Gene Expression Profiling Interactive Analysis) web-based platform serves as a centralized source of GTEx and TCGA gene expression data [35]. Using the Correlation Analysis function from GEPIA2, Pearson’s correlation coefficient was determined for 33 TCGA tumor tissues, and we identified several cancers where CAPN4 and Bsg expression was positively correlated (Fig. 7). Statistical significance of each correlation’s strength was also determined by GEPIA2 using a t-test for Pearson’s correlation coefficient.

## RESULTS

### Basigin is a Binding Partner of Calpain 4

To further elucidate the mechanism by which calpain4 produces traction force, the complete Capn4 gene was inserted into the plasmids pCWX200 and pLexA and used as bait to identify binding partners for the calpain 4 protein in a yeast two-hybrid assay. Sequencing identified basigin as one candidate binding partner for calpain 4. This direct binding between calpain 4 and basigin was confirmed by co-immunoprecipitation (Fig. 1 *B*). The direct binding between calpain 4 and basigin suggests the possibility that basigin is involved in the pathway for the generation of traction forces regulated by calpain 4.

**FIGURE 1.**
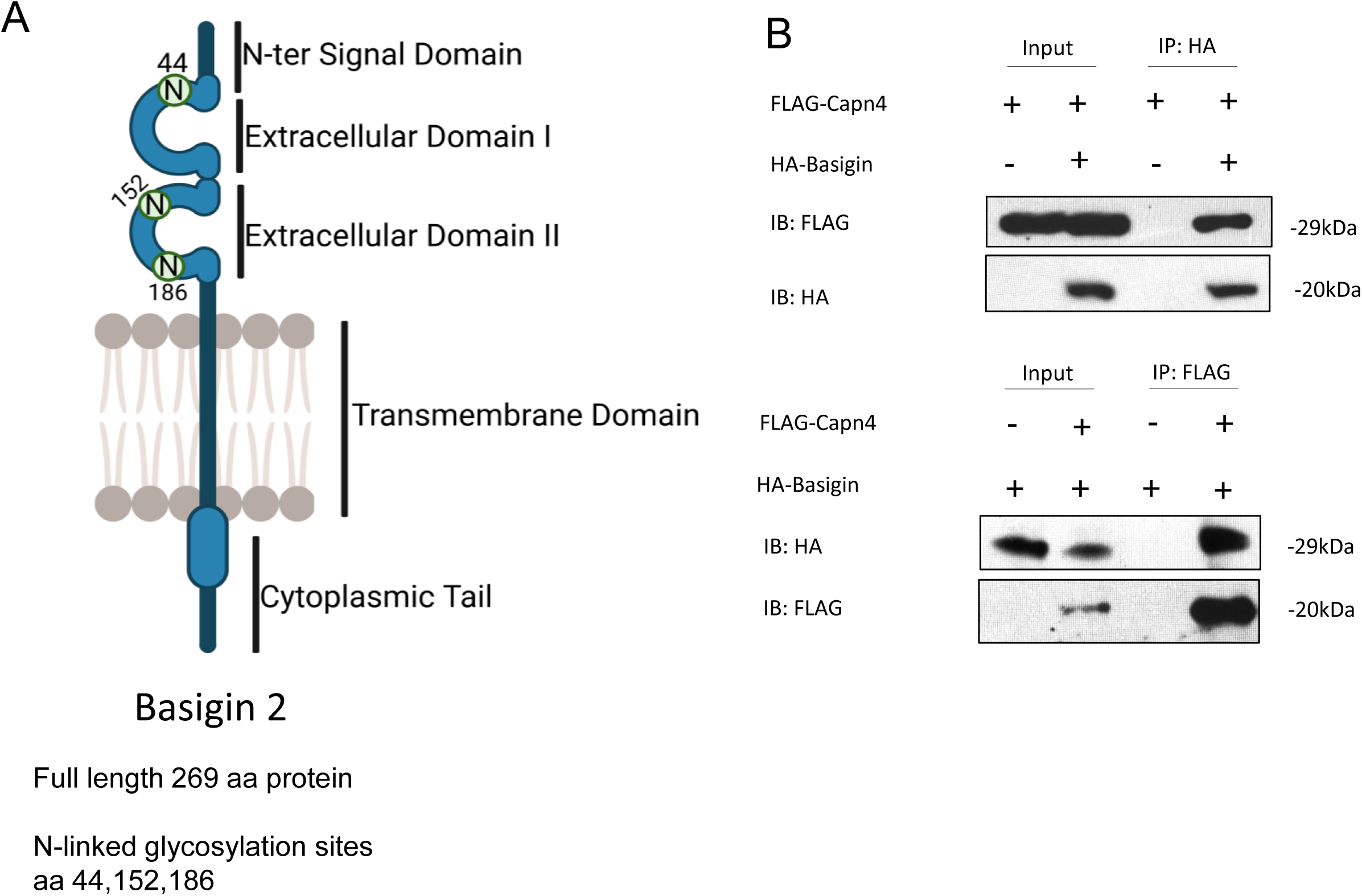
Co-immunoprecipitation to confirm the direct binding between calpain4 and basigin proteins. *A*. Illustration of the transmembrane protein basigin 2 (modified Biorender illustration). *B.* The entire gene of *Capn4* was inserted into a pFLAG-CMV vector, and the Bsg gene, lacking a 100 amino acid sequence for the N-terminal, was inserted into a pCDNA3 vector together with an HA sequence. Both plasmids were transfected into 293T cells. Pull-down assays were performed using either FLAG or HA antibody to confirm the direct binding between calpain 4 and basigin.

When comparing the expression level of basigin in both MEFs and *Capn4-/-* cells, we observed that basigin was expressed at a reduced level in *Capn4-/-* cells compared to MEFs (Fig. 2 *A*), as supported by quantification (Fig. 2 *B*) (Supporting information 1A). Basigin is known to be glycosylated on three asparagines (Fig. 1. *A*). The glycosylation state can result in multiple bands and is referred to as high and low glycosylation states. Glycosylation can be demonstrated by enzyme digestion with PNGase (Supporting Information 1*B*). Interestingly, in the absence of calpain4, a lower glycosylation state of Basigin is observed in MEF cells (Fig. 2*A*, *B*); however, the significance of these glycosylation states will be explored in the discussion.

**FIGURE 2.**
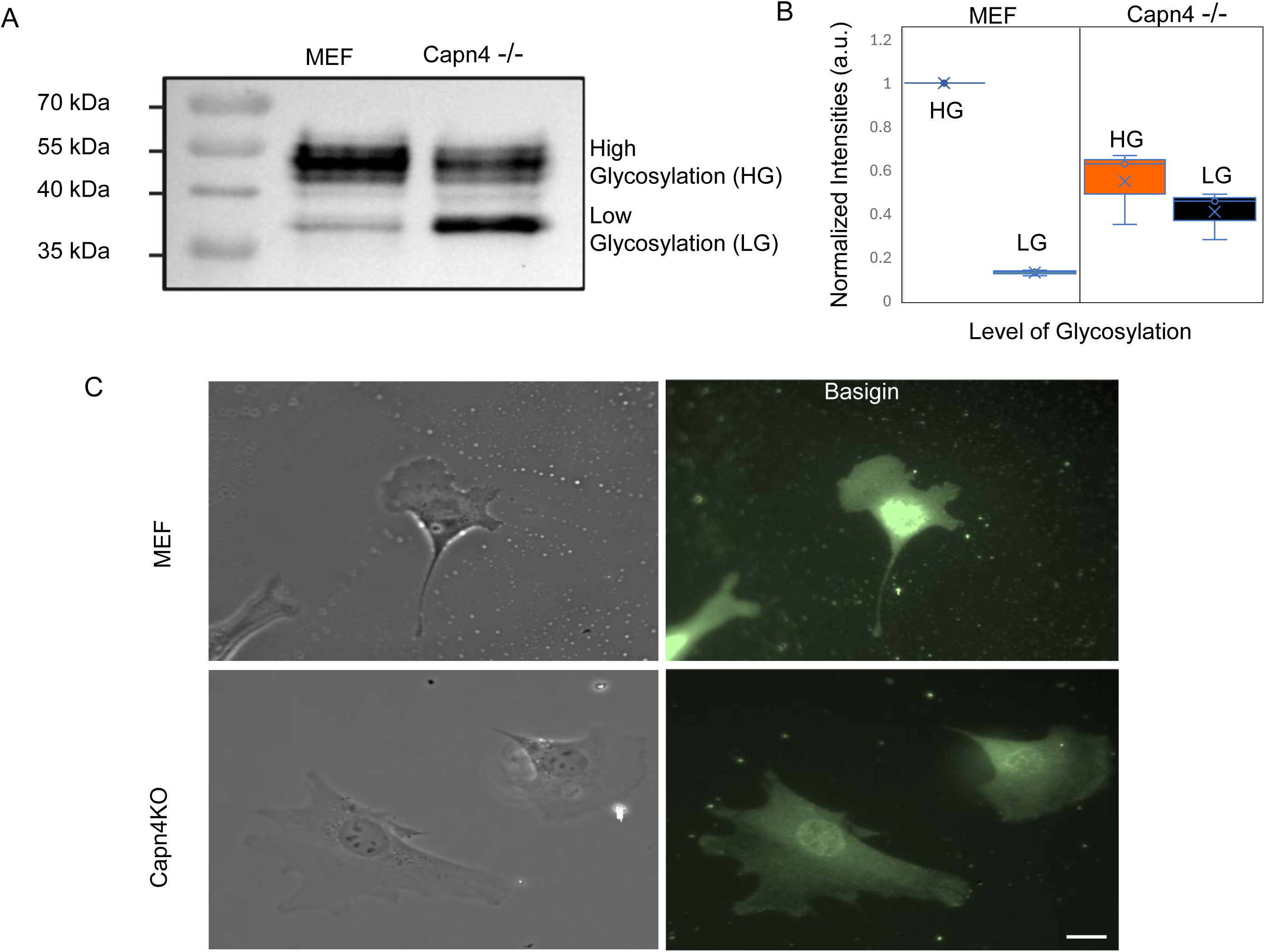
Basigin expression level is reduced in *Capn4-/-* cells. *A.* Two bands of 50 and 37 kDa were detected by western blot probed with monoclonal anti-basigin. The bands represent high and low levels of basigin glycosylation, respectively. *B*. Intensities normalized to total protein (Supplemental 1*A)* and glycosylation confirmed (Supplemental 1B)*. C.* Localization of basigin in MEFs and *Capn4-/-* cells. Cells were seeded onto fibronectin-coated coverslips. ImageJ calculated the corrected total cell fluorescence (CTCF) for each cell line. *Capn4^−/–^* cells exhibit significantly reduced levels of CTCF when compared to MEF. 12 cells were used for each cell line for calculation. Data for *A.* and *B.* represent three independent trials. (Mag. bar = 20μm).

To observe the expression pattern of the basigin protein and investigate potential co-localization with calpain 4, immunofluorescence was performed in both MEFs and Capn4-/-cells using a basigin antibody. Basigin is predominantly localized perinuclear and diffusely at the cell edge (Fig. 2*C*). Basigin does not localize to the periphery of *Capn4-/-* cells. However, this result is likely due to the thinness of the lamellipodia in Capn4-/-cells, as previously described [14]. Combined, we find evidence of direct interaction between basigin and calpain 4 via two-hybrid and co-immunoprecipitation, as well as differences in glycosylation and cellular immunofluorescence of basigin in capn4-/-cells.

### Knockdown of Basigin Resulted in Defects in Traction Force and Adhesion Strength in MEFs

Previous studies in cell migration revealed a deficiency in traction forces in *Capn4^−/–^* cells compared to wild-type MEF cells, while inhibition of *Capn1* or *Capn2* or the overexpression of calpastatin did not affect traction forces [14, 15]. Given the direct binding of basigin and calpain 4, we knocked down basigin and asked if it would impact the production of traction force. Basigin was knocked down by nucleofection of siRNA in MEF cells, resulting in a high efficiency (95%) of inhibition after 36 hours, as demonstrated by western blots and quantification (Supporting information 2).

To measure the traction stress generated by the basigin knockdown, cells were seeded onto flexible polyacrylamide substrate covalently coated with fibronectin and subjected to analysis by traction force microscopy (TFM) [33]. Traction stress in each cell line was measured (Fig. 3 *A, B, C*). As expected, *Capn4^−/–^* cells produced significantly less traction stress (avg. 1.5 kPa) compared to wild-type MEFs (avg. 2.69kPa, *p*=0.03) and MEFs transfected with control siRNA (avg. 2.91kPa, *p*=0.04). Furthermore, silencing basigin in MEFs also significantly reduced the magnitude of traction forces to 1.63 kPa (*p*=0.04) (Fig. 3. *A, B, C*). These results demonstrate that silencing basigin leads to a deficiency in the production of traction stress comparable to that obtained from the disruption of calpain 4.

**FIGURE 3.**
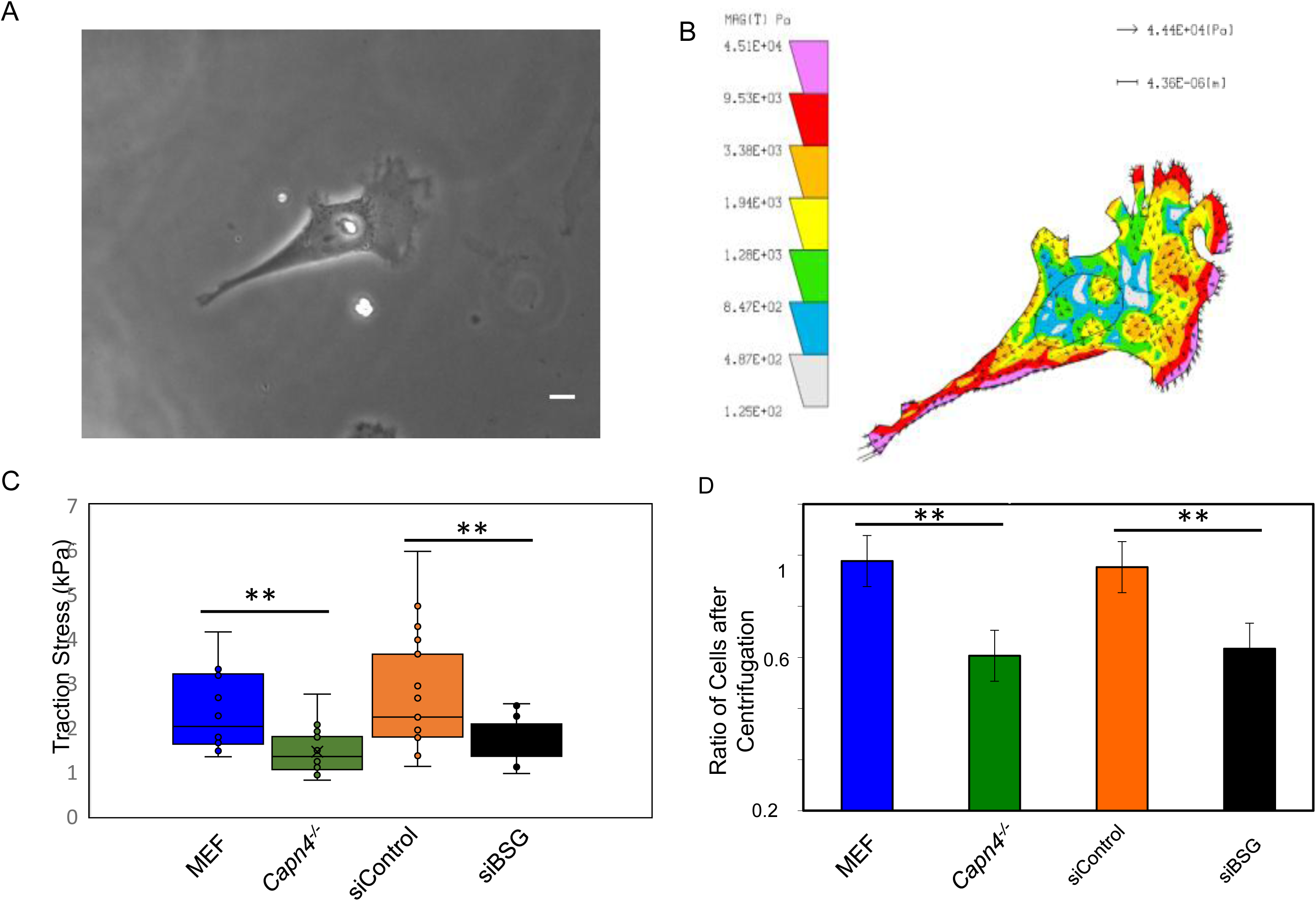
Basigin traction stress and adhesion strength mimics Capn4 -/-. *A.* A phase image representing an MEF cell nucleofected with siBSG and seeded on fibronectin-coated polyacrylamide substrates overnight before TFM. *B.* A color map of the traction stress of the cell in the image of panel *A.* The map depicts warm colors as high stress and cool colors as low stress. The arrows indicate the magnitude, location, and direction of the stress. The strongest stress is found at the leading edge, consistent with previously reported capn4-/-TFM. *C.* A graph representing the average traction stress exerted by each cell line onto the substrate. Traction stress of basigin knock-down MEFs was compared with MEFs, *Capn4^−/–^* cells, and MEFs transfected with control siRNA. When basigin was silenced by siRNA, the traction stress level was significantly reduced, as well as in *Capn4^−/–^* cells, compared to MEFs and MEFs transfected with control siRNA. 14 MEFs, 22 *Capn4^−/–^*cells, 21 MEFs transfected with control siRNA, and 21 basigin knockdown MEFs were used for traction force calculation. *D.* A bar graph representing the adhesion strength of cells by calculating the percentage of the number of cells that remained adhered onto the substrates after centrifugation and compared to MEFs, *Capn4^−/–^* cells exhibited significantly reduced adhesion strength (*p*=0.02). When basigin was silenced through siRNA in MEFs, a reduction of adhesion strength was also observed (*p*=0.02). Three independent experiments are represented. Students’ t-test performed statistical analysis.

To test the adhesion strength of focal adhesions to the substrates, we performed a centrifugation assay with the same set of cell lines as above [14, 15, 34]. Cells were seeded onto fibronectin-coated flexible polyacrylamide substrates and allowed to adhere for 30 minutes at 37°C before centrifugation. The number of cells adhered to the substrates was counted before and after centrifugation. We found that approximately 61% of Capn4–/– cells remained adhered to the substrates after centrifugation, compared to 98% of MEFs that remained adhered (Fig. 3*D*). Similarly, silencing basigin resulted in only approximately 63% of cells remaining adhered on the substrates (Fig. 3 *D*). In comparison, 95% percent of MEFs treated with control siRNA remained adhered after centrifugation (Fig. 3 *D*). These results suggest that basigin contributes to the adhesion strength of focal adhesions, in addition to regulating the production of traction force, consistent with the behavior of *Capn4-/-*.

### Mechanical Tension is not affected by the Knockdown of Basigin

Cells can sense mechanical information from the environment, including matrix elasticity, localized mechanical forces, and topography [36–40]. These physical signals are transmitted outside-in and lead to changes in the cytoskeletal networks, interaction with the extracellular matrix (ECM), cellular force production, differentiation, growth, and apoptosis [36, 37, 39–41]. Our previous studies found that MEFs respond to localized mechanical stimulation by changing their migration trajectory or rounding up. However, when cells deficient in either Calpain 1, 2, or 4 are tested in this assay, they continue to migrate along the same trajectory, indicating they are insensitive to localized stimuli [14, 15]. In another assay comparing the ability of cells to spread on substrates of different stiffness, the results indicate that MEFs can sense stiffness by spreading more completely on stiff substrates compared to soft substrates [15, 42]. Surprisingly, MEF cells deficient in any calpain 1, 2, or 4 are still able to sense the stiffness difference and spread differently on hard and soft substrates [14, 15]. Traction forces were believed to not only function as the driving force for cell migration but also play equal roles in sensing the physical environment [36]. As our study indicated that silencing basigin in MEFs significantly affected the generation of traction forces, we tested basigin knockdown cells for their ability to sense changes in both mechanosensing assays.

Basigin knockdown cells were tested for their ability to respond to localized mechanical stimuli. In the assay, cells were seeded onto flexible polyacrylamide substrates coated with fibronectin, and a blunted needle was gently pushed onto the substrate against the direction of migration. As expected, 88% of MEF cells responded to the pushing force by avoiding it while only 14% of *Capn4^−/–^* cells responded (Fig. 4 *A, B*). Like MEF controls, 84% of MEF cells transfected with control siRNA reacted to the localized pushing force (Fig. 4 *A, B*). When basigin was silenced in MEFs, 84% of cells still responded to localized force (Fig. 4*A, B*). Our results indicate that basigin does not play a role in sensing localized mechanical stimulus.

**FIGURE 4.**
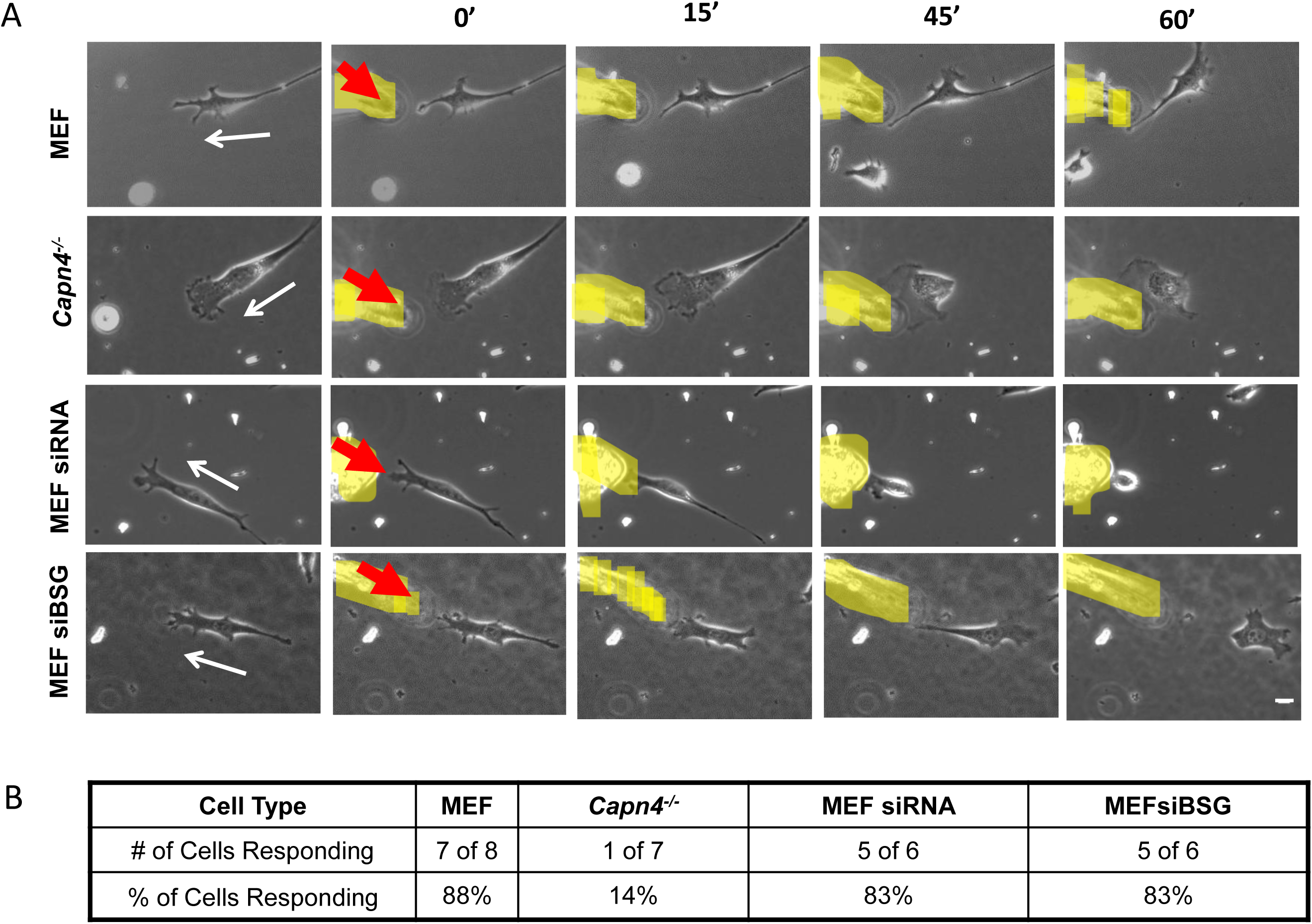
Silencing basigin does not affect the ability of cells to sense changes in the mechanical environment. *A*. Representative time-lapse images illustrate the responses of cells to a localized stimulus, including MEFs, *Capn4^−/–^*cells, MEFs transfected with control siRNA, and basigin knockdown MEFs. The thin white arrows in the first column denote the original cell migration direction; the bold red arrows in the second column denote the direction in which the blunted microneedle pushes on the substrate, thereby making the substrate softer in front of the migrating cell. The blunted microneedle is highlighted in yellow. Cells were seeded onto flexible polyacrylamide substrates that were covalently coated with fibronectin and allowed to attach overnight. A blunted needle was inserted into the substrate along the migration path, and the responses of cells were recorded for each cell line (magnification bar = 10 μm) and summarized in *B*. The table indicates the percentage of cells of each cell line that respond to the localized change in the mechanical environment. The number of cells and percentage response for each cell line were summarized in the table. If a cell avoids the pushing force or rounds up, it is sensing and responding to the change; if a cell continues to migrate toward the pushing force, it is not sensing the localized change in stiffness.

To test if cells deficient in basigin can sense a more global homeostatic change in the stiffness of the environment, cells were seeded onto hard and soft flexible polyacrylamide substrates coated with fibronectin and allowed to adhere overnight. Cells were scored for a rounded phenotype. As expected, when seeded on rigid substrates, 85% of MEFs spread normally on hard substrates, as well as 91% of *Capn4^−/–^* cells. Meanwhile, 86% of MEFs treated with control non-target siRNA and 90% of MEFs treated with basigin-targeting siRNA also spread normally (Fig. 5*A, B*). In contrast, only 47% of MEFs, 85% of *Capn4^−/–^* cells, 37% of MEFs treated with control non-target siRNA, and 44% of MEFs treated with basigin targeting siRNA spread well when seeded on soft substrates (Fig. 5 *A, B*). The significant decrease in the number of cells spreading normally on substrates of different stiffness indicated that basigin is not involved in sensing the stiffness of the environment. Combined with the cellular responses to a localized stimulus, unlike calpain-4, basigin is not significant to the mechanosensing process.

**FIGURE 5.**
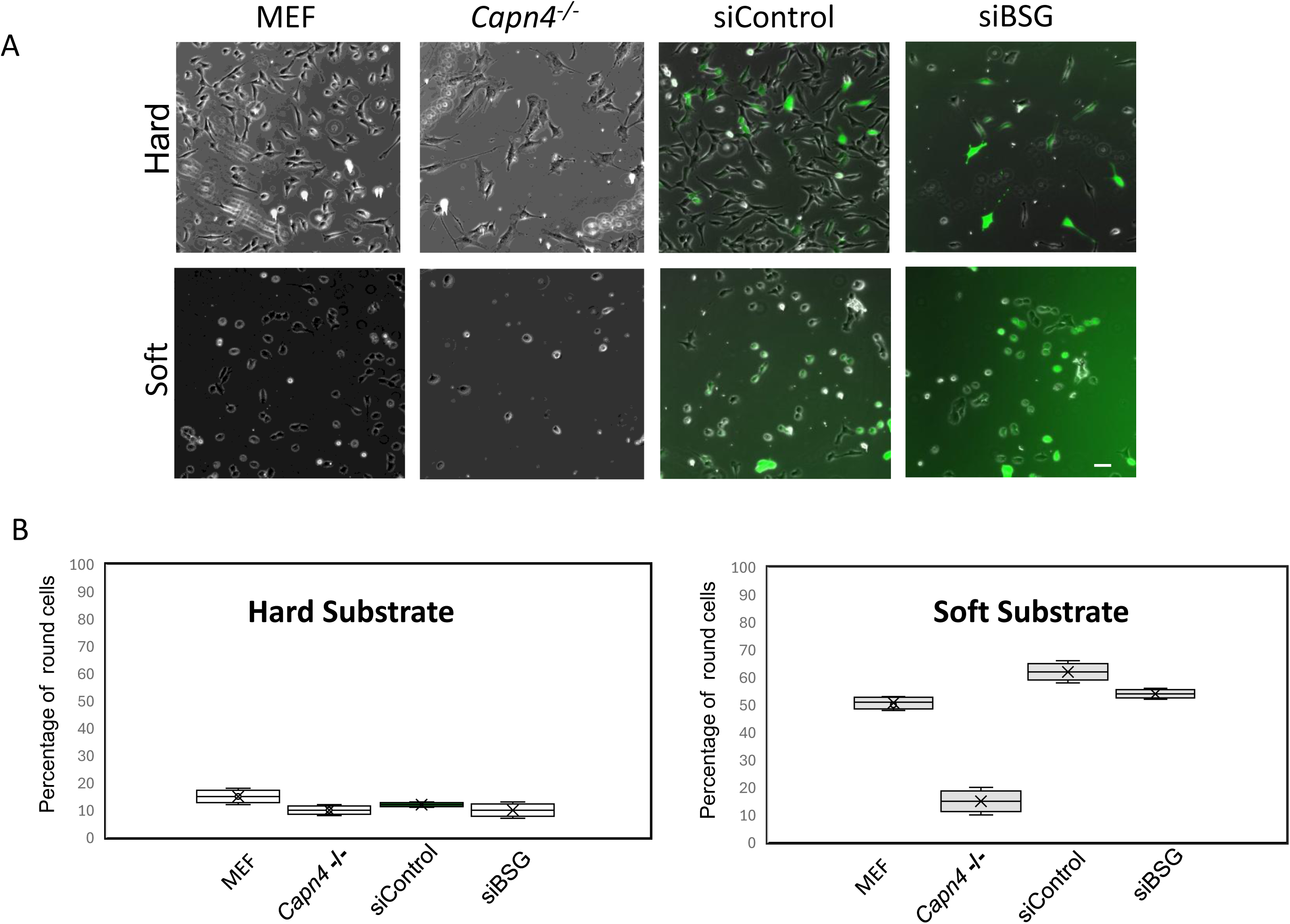
Basigin-deficient cells can sense a soft environment, but capn4-/-cells do not. *A.* Images were taken with a 10X lens for each cell line after they were seeded on both hard (5%/0.1% Acryl/Bis) and soft (5%/0.04% Acryl/Bis) substrates and allowed to adhere overnight. The number of round cells was then counted for each line as observed visually. *B.* The average cell counts for each line were graphed; six random fields were counted for each cell line. Statistical analysis was performed by Student’s t-test. * indicates *p*<0.05.

### Knockdown of Basigin Does not Affect Cell Migration Speed and Migration Persistence

The speed and persistence of cell migration are affected by multiple factors, including dimension, matrix stiffness, cell–cell and cell–matrix adhesion, traction forces, cytoskeletal polarity, and the capacity to degrade ECM by proteolytic enzymes [43, 44]. Previous observations indicate that Capn4–/– cells have a reduced migration speed compared to MEF cells [11, 14] when cultured on fibronectin-coated coverslips; however, this difference is not observed on fibronectin-coated polyacrylamide substrates [15]. To learn whether inhibition of basigin impacted cell migration, basigin knockdown cells were seeded onto fibronectin-coated polyacrylamide substrates and observed for 2 hours to track the locomotion of the nuclei. Consistent with previous studies, *Capn4^−/–^* cells migrated at linear speeds of (0.45 μm/min) compared to MEFs (0.48 μm/min, *p=*0.60). When basigin was silenced, an insignificant difference in linear speed was also observed, such that MEFs+siBSG migrated at 0.47 μm/min, p=0.65 on fibronectin-coated polyacrylamide substrates (Fig. 5, *A*). As with migration speed, silencing basigin did not affect migration persistence and was similar in all lines (Fig. 5, *B*). These results demonstrate that the mechanism of calpain 4, which also involves basigin and regulates traction generation, does not impact linear migration speed or persistence.

### Correlation of Gene Expression of CAPN4 and BSG in Various Human Tumors

In previous studies, both BSG and CAPN4 were independently implicated in tumor progression [45, 46]. Based on this knowledge, we investigated whether their expression was correlated across an array of tumors. Using the GEPIA2 platform, which integrates gene expression data from GTEx and TCGA databases, Pearson’s correlation coefficients were determined to measure the correlative strength of CAPN4 and BSG expression in several tumors (Fig. 7*A*) [35]. Amongst these tumors, R-values ranged from moderate (R > 0.30) to strong (R > 0.50), and all correlations were considered positive and statistically significant (p < 0.001). The strongest correlations were observed in Uveal Melanoma (R = 0.75, p = 1.1 × 10^ (-15)) and Glioblastoma (R = 0.60, p = 0) tumors, indicating the co-expression of BSG and CAPN4 in these malignancies. Furthermore, the most moderate correlations were observed in Brain Lower Grade Glioma (R = 0.4, p = 0) and Kidney Renal Papillary Cell Carcinoma (R = 0.42, p = 1.6E-013) tumors (Figure 7 *B*). Of particular interest is the shared neural origin and aggressive nature of Uveal Melanoma and Glioblastoma, suggesting that the co-expression of BSG and CAPN4 may contribute more significantly to tumor aggressiveness in cancers of neural origin. Collectively, these results indicate that, although the strength of correlation varies, the broader presence of CAPN4 and BSG correlative expression across diverse tumor tissues may be associated with enhanced tumor progression and cancer progression.

## DISCUSSION

Our lab has previously discovered that Calpains participate in both the generation of traction stress and sensing localized stimuli in MEF cells [14]. We found that both large and small subunits of calpain holoenzymes are required for cells to sense localized stimuli normally, while only the small subunit is necessary to create traction stress; however, there is no effect on the production of traction stress when the large catalytic subunits are silenced [14]. This suggests that the calpain small subunit functions independently of the proteolytic large subunits of calpain in the regulation of traction forces, while all subunits are implicated in mechanosensing. In this study, we sought to gain insight into how this mechanism works by identifying direct binding partners of Calpain4.

We have discovered a previously unknown physical interaction between Basigin and Calpain-4, as determined by two-hybrid and co-immunoprecipitation assays. We also looked for co-localization by immunofluorescence. We were not surprised to find a lack of co-localization between basigin and calpain4 via conventional immunofluorescence, given that calpain4 localization is diffuse on its own. What was worth noting was the diminished signals for basigin location at the periphery of *Capn4-/-* cells. Basigin is known to have a diverse localization pattern, often dependent on the cell type. As expected, it frequently localizes to the plasma membrane; interestingly, it can also be found in focal adhesions, amongst other cellular locations. The high expression level and multiple functions of basigin in varied types of cells likely explain the lack of localization of basigin in our study (Fig. 2*C*) [18–22].

Studies of basigin in cell migration have focused on its function in tumor cell motility and invasion. It has been reported that basigin expression is commonly elevated in most types of tumor cells and is one of the most highly expressed proteins in disseminated cancer cells [24]. A high level of basigin expression on the surface of tumor cells induces an increased level of MMP activity in both stromal cells and the tumor cells themselves [17, 47, 48]. Elevated MMP activity degrades the ECM and alters ECM turnover dynamics, potentially leading to tumor cell motility and invasion [24]. Consistent with these observations, our current study found that inhibiting basigin expression through siRNA in wild-type MEFs results in reduced traction force and adhesion strength (Fig. 3C, D); however, cells had normal migration speed (Fig. 6*A*), mirroring the calpain4 knockout and suggesting that calpain 4 functions with basigin in this pathway. However, basigin affects numerous targets in addition to MMPs [17, 24]. Other proteins are likely involved in regulating traction stress through this pathway. Further studies are needed to identify the components that function downstream of this signaling pathway.

**FIGURE 6.**
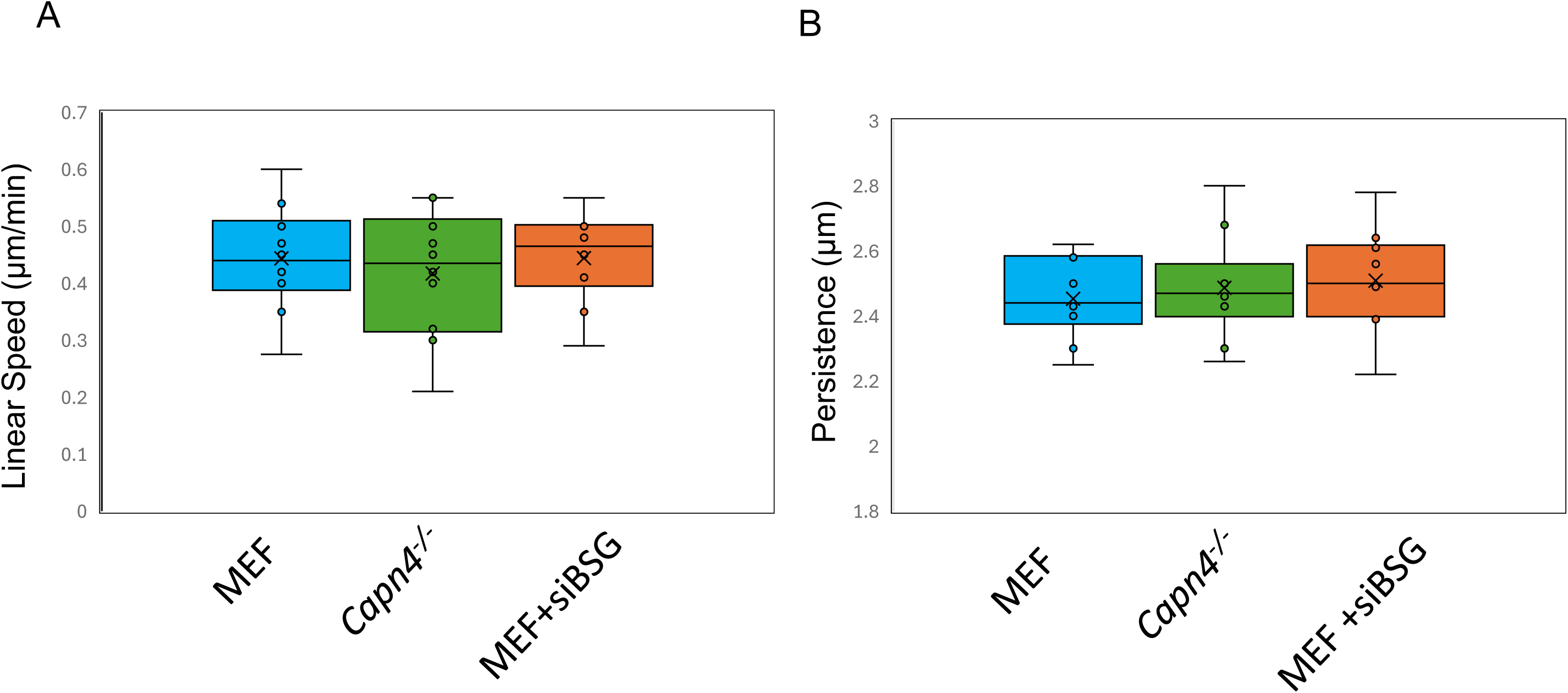
Cell migration and persistence are not affected by siBSG and capn4-/-cells when measured on fibronectin-coated polyacrylamide gels. *A*. The bar graph represents the average migration speed of MEFs, *Capn4^−/–^* cells, and MEFs with basigin knocked down on polyacrylamide gels covalently coated with fibronectin. *B*. The persistence of migration was calculated for each cell line. No significant difference in persistence was observed among the three cell lines. 18 MEF cells, 15 *Capn4^−/–^* cells, and 15 basigin knockdown MEF cells were used for calculation in A and B. Statistical analysis was performed by Student’s t-test.

**Figure 7.**
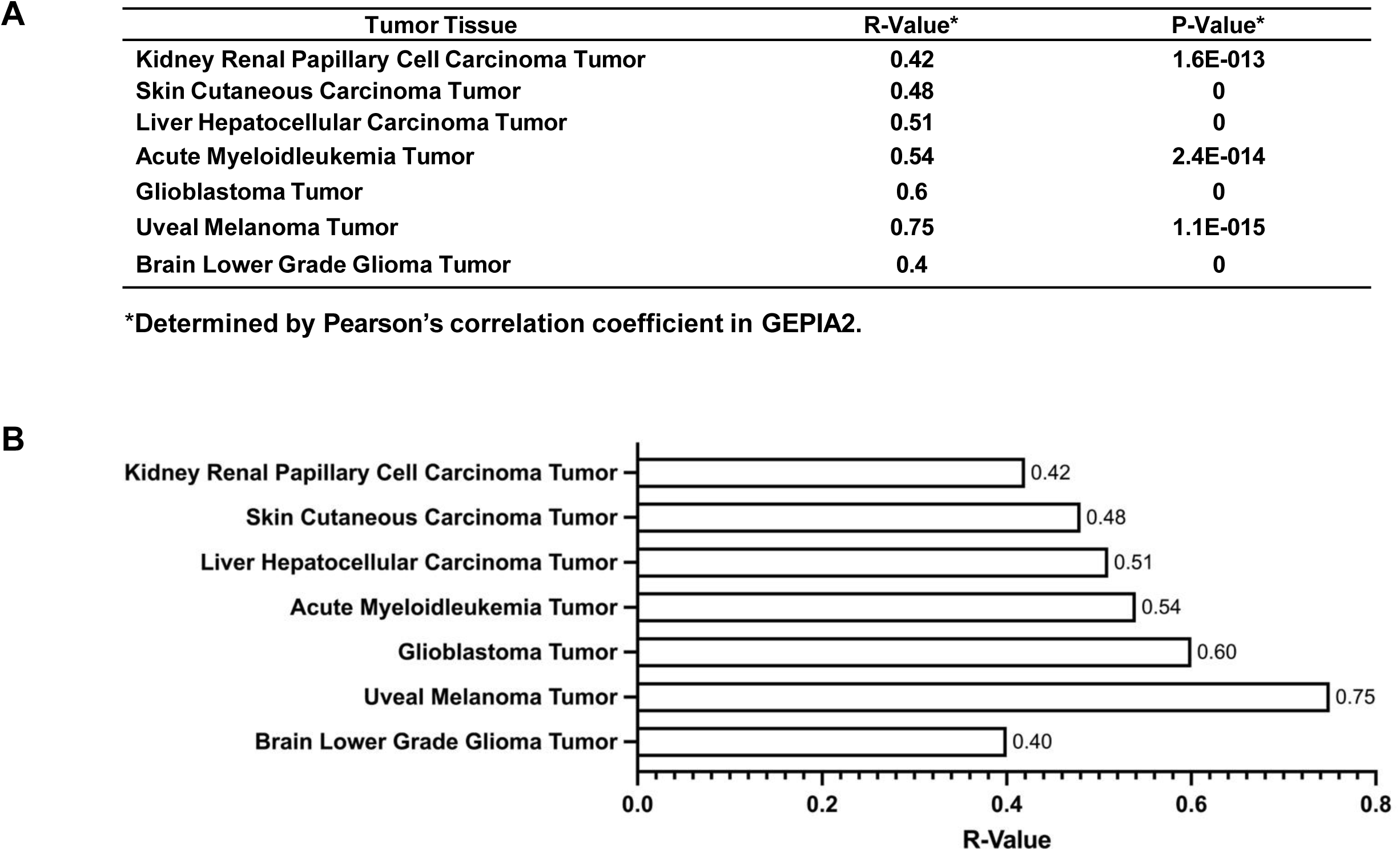
R and P-Values for CAPN4 and BSG Expression Correlation in Various Tumor Tissues. *A.* Pearson’s correlation analysis was performed using GEPIA2 (Tang et al., 2019), which enabled the modeling of CAPN4 and BSG expression across seven tumor tissues sourced in the TCGA database. From the Correlational Analysis function of GEPIA2, R and p-values, determined by Pearson’s correlation coefficient, for each of the respective tissues were collected and modeled in a table. The R-value is specific to the strength and direction of the correlation between CAPN4 and BSG expression, where a value greater than 0.5 indicates a strong, positive relationship, and a value between 0.3 and 0.5 demonstrates a moderate, positive relationship. Further, the p-value validates statistical significance. All reported correlations were statistically significant (p < 0.001). *B.* A bar graph demonstrating the R-values of the correlation between CAPN4 and BSG expression in seven tumors of interest was generated.

Previous research indicates a deficiency in traction force in Capn4-/-cells (Fig. 3*C*). Further, these *Capn4-/-* cells do not respond to localized stimuli and do not sense the stiffness of substrates normally (Fig. *4 A.* and Fig *5A*), suggesting that calpain4 has provided a means to separate traction force and mechanosensing spatially and temporally. Moreover, MEFs in which basigin is silenced can respond to the localized stimuli and sense the stiffness of substrates normally (Fig. 4 *A* and Fig. 5*D*), suggesting a function of basigin only in the production of traction forces but not in mechanosensing. Previous studies indicate that substrate rigidity sensing is driven by traction forces at the cells’ leading edge, as localized softening of the frontal region results in retraction of the cell, reversal of cell polarity, or cell immobilization [49]. Our conclusion does not contradict this observation, as we only measured the overall level of traction stress within a cell, without measuring localized areas of the cell.

The difference in the glycosylation state of Basigin in calpain-4-deficient cells was unexpected, though not without precedent. It is well documented that glycosylation is sensitive to various culture conditions, including temperature, pH, media composition, and cell passage [50]. However, our observation of a lower glycosylation state of Basigin in *Capn4*⁻/⁻ cells remains consistent. There is a precedent that the degree of Basigin’s glycosylation can regulate its function [51]. In our study, we speculate that it modulates the binding of Galectin-3, a lectin that facilitates the clustering and activation of both Basigin and integrin β1 [52]. We have previously shown that Galectin-3 secretion requires calpain 4, and that exogenous Galectin-3 can rescue the traction force defect [15]. Although this mechanism is beyond the scope of the present study, it is currently under further investigation.

Basigin has been reported to regulate integrin β1 via direct interaction; however, the literature presents conflicting evidence. In *Drosophila* perineurial glia, Basigin has been described as a negative regulator, while in other systems, it functions as a positive regulator [53, 54]. Given that Galectin-3 promotes the clustering of Basigin and integrin β1, and that Galectin-3-secretion-deficient mutants (Gal3 Y107F) exhibit reduced levels of active integrin β1, it is plausible that Basigin serves as an activator of integrin β1 in this context [15]. The interplay between Basigin’s glycosylation state, its lateral organization at the membrane, Galectin-3 binding, and integrin β1 activation may constitute a dynamic regulatory axis underlying the generation of traction forces.

The link between calpain4 and basigin is intriguing and has left us to question how often the expression levels correlate in cancers. As discussed, basigin is known to affect the progression of cancer and to act as a stimulator of ECM degradation. Still, until now, the possibility of calpain 4 being involved in cancer progression along with basigin was unknown. Furthermore, the high correlation in neural cancers presents an avenue for new research and potential therapeutic targeting opportunities.

In summary, we have identified a direct binding partner for Calpain4 in Basigin. Further, we have determined that Basigin and Calpain4 contribute to the regulation of traction force and the strength of substrate adhesion. However, Basigin and Calpain4 are not implicated in mechanosensing and do not respond to localized stimuli and homeostatic tension. Together, these results implicate basigin in an unknown pathway in which calpain 4 is involved in regulating traction force independently of the catalytic large subunits of calpain 1 and 2.

## Acknowledgements

The authors would like to acknowledge Dr. Alexander Gasparski for writing and technical assistance. Portions of this work were developed from the dissertation of Dr. Bingqing Hao. We would also like to thank Dr. Ochrietor for the basigin plasmid and Dr. Zhang for the contribution of the GEPIA2 platform. Funding for this project was initially provided to KAB through NIH R01-GM084248 and start-up funds from Wayne State University.

## Notes

### Competing Interest Statement

The authors have declared no competing interest.

### Summary of Updates

We have found a glycosylation effect of basigin and included a gene expression correlation analysis, and replaced some of the western data. In addition, many of the bar graphs have been converted to box and whisker plots to display the spread in data more accurately. A significant rewrite of the discussion has been done.

## References

1 Lauffenburger, D. A. and Horwitz, A. F. (1996) Cell Migration: A Physically Integrated Molecular Process. Cell 84, 359–369 10.1016/S0092-8674(00)81280-5

2 Friedl, P. and Alexander, S. (2011) Cancer Invasion and the Microenvironment: Plasticity and Reciprocity. Cell 147, 992–1009 10.1016/j.cell.2011.11.016

3 Walters, N. J. and Gentleman, E. (2015) Evolving insights in cell–matrix interactions: Elucidating how non-soluble properties of the extracellular niche direct stem cell fate. Acta Biomater. 11, 3–16 10.1016/j.actbio.2014.09.038

4 Whelan, D., Caplice, N. M. and Clover, A. J. P. (2014) Fibrin as a delivery system in wound healing tissue engineering applications. J. Controlled Release 196, 1–8 10.1016/j.jconrel.2014.09.023

5 Fouchard, J., Bimbard, C., Bufi, N., Durand-Smet, P., Proag, A., Richert, A., et al. (2014) Three-dimensional cell body shape dictates the onset of traction force generation and growth of focal adhesions. Proc. Natl. Acad. Sci. 111, 13075–13080 10.1073/pnas.1411785111

6 Goldmann, W. H. (2014) Mechanosensation. In Progress in Molecular Biology and Translational Science, pp 75–102, Elsevier 10.1016/B978-0-12-394624-9.00004-X

7 Pasapera, A. M., Plotnikov, S. V., Fischer, R. S., Case, L. B., Egelhoff, T. T. and Waterman, C. M. (2015) Rac1-Dependent Phosphorylation and Focal Adhesion Recruitment of Myosin IIA Regulates Migration and Mechanosensing. Curr. Biol. 25, 175–186 10.1016/j.cub.2014.11.043

8 Ridley, A. J., Schwartz, M. A., Burridge, K., Firtel, R. A., Ginsberg, M. H., Borisy, G., et al. (2003) Cell Migration: Integrating Signals from Front to Back. Science 302, 1704–1709 10.1126/science.1092053

9 Beckerle, M. C., Burridge, K., DeMartino, G. N. and Croall, D. E. (1987) Colocalization of calcium-dependent protease II and one of its substrates at sites of cell adhesion. Cell 51, 569–577 10.1016/0092-8674(87)90126-7

10 Bhatt, A., Kaverina, I., Otey, C. and Huttenlocher, A. (2002) Regulation of focal complex composition and disassembly by the calcium-dependent protease calpain. J. Cell Sci. 115, 3415–3425 10.1242/jcs.115.17.3415

11 Dourdin, N., Bhatt, A. K., Dutt, P., Greer, P. A., Arthur, J. S. C., Elce, J. S., et al. (2001) Reduced Cell Migration and Disruption of the Actin Cytoskeleton in Calpain-deficient Embryonic Fibroblasts. J. Biol. Chem. 276, 48382–48388 10.1074/jbc.M108893200

12 Goll, D. E., Thompson, V. F., Li, H., Wei, W. and Cong, J. (2003) The Calpain System. Physiol. Rev. 83, 731–801 10.1152/physrev.00029.2002

13 Xu, L. and Deng, X. (2006) Suppression of Cancer Cell Migration and Invasion by Protein Phosphatase 2A through Dephosphorylation of μ- and m-Calpains. J. Biol. Chem. 281, 35567–35575 10.1074/jbc.M607702200

14 Undyala, V. V., Dembo, M., Cembrola, K., Perrin, B. J., Huttenlocher, A., Elce, J. S., et al. (2008) The calpain small subunit regulates cell-substrate mechanical interactions during fibroblast migration. J. Cell Sci. 121, 3581–3588 10.1242/jcs.036152

15 Jang, I., Menon, S., Indra, I., Basith, R. and Beningo, K. A. (2024) Calpain Small Subunit Mediated Secretion of Galectin-3 Regulates Traction Stress. Biomedicines 12, 1247 10.3390/biomedicines12061247

16 Muramatsu, T. and Miyauchi, T. (2003) Basigin (CD147): a multifunctional transmembrane protein involved in reproduction, neural function, inflammation and tumor invasion. Histol. Histopathol. 981–987 10.14670/HH-18.981

17 Gabison, E. E., Hoang-Xuan, T., Mauviel, A. and Menashi, S. (2005) EMMPRIN/CD147, an MMP modulator in cancer, development and tissue repair. Biochimie 87, 361–368 10.1016/j.biochi.2004.09.023

18 Chen, H., Fok, K. L., Yu, S., Jiang, J., Chen, Z., Gui, Y., et al. (2011) CD147 is required for matrix metalloproteinases-2 production and germ cell migration during spermatogenesis. Mol. Hum. Reprod. 17, 405–414 10.1093/molehr/gar013

19 Igakura, T., Kadomatsu, K., Kaname, T., Muramatsu, H., Fan, Q.-W., Miyauchi, T., et al. (1998) A Null Mutation in Basigin, an Immunoglobulin Superfamily Member, Indicates Its Important Roles in Peri-implantation Development and Spermatogenesis. Dev. Biol. 194, 152–165 10.1006/dbio.1997.8819

20 Saxena, D., Oh-Oka, T., Kadomatsu, K., Muramatsu, T. and Toshimori, K. (2002) Behaviour of a sperm surface transmembrane glycoprotein basigin during epididymal maturation and its role in fertilization in mice. Reproduction 123, 435–444 10.1530/rep.0.1230435

21 Igakura, T., Kadomatsu, K., Taguchi, O., Muramatsu, H., Kaname, T., Miyauchi, T., et al. (1996) Roles of Basigin, a Member of the Immunoglobulin Superfamily, in Behavior as to an Irritating Odor, Lymphocyte Response, and Blood–Brain Barrier. Biochem. Biophys. Res. Commun. 224, 33–36 10.1006/bbrc.1996.0980

22 Hori, K., Katayama, N., Kachi, S., Kondo, M., Kadomatsu, K., Usukura, J., et al. (2000) Retinal Dysfunction in Basigin Deficiency. Invest. Ophthalmol. Vis. Sci. 41, 3128–3133

23 Liu, F., Cui, L., Zhang, Y., Chen, L., Wang, Y., Fan, Y., et al. (2010) Expression of HAb18G is associated with tumor progression and prognosis of breast carcinoma. Breast Cancer Res. Treat. 124, 677–688 10.1007/s10549-010-0790-6

24 Xiong, L., Edwards, C. and Zhou, L. (2014) The Biological Function and Clinical Utilization of CD147 in Human Diseases: A Review of the Current Scientific Literature. Int. J. Mol. Sci. 15, 17411–17441 10.3390/ijms151017411

25 Yang, M., Yuan, Y., Zhang, H., Yan, M., Wang, S., Feng, F., et al. (2013) Prognostic significance of CD147 in patients with glioblastoma. J. Neurooncol. 115, 19–26 10.1007/s11060-013-1207-2

26 Biswas, C., Zhang, Y., DeCastro, R., Guo, H., Nakamura, T., Kataoka, H., et al. (1995) The human tumor cell-derived collagenase stimulatory factor (renamed EMMPRIN) is a member of the immunoglobulin superfamily. Cancer Res. 55, 434–439

27 Kanekura, T., Chen, X. and Kanzaki, T. (2002) Basigin (cd147) is expressed on melanoma cells and induces tumor cell invasion by stimulating production of matrix metalloproteinases by fibroblasts. Int. J. Cancer 99, 520–528 10.1002/ijc.10390

28 Li, Y., Wu, J., Song, F., Tang, J., Wang, S.-J., Yu, X.-L., et al. (2012) Extracellular Membrane-proximal Domain of HAb18G/CD147 Binds to Metal Ion-dependent Adhesion Site (MIDAS) Motif of Integrin β1 to Modulate Malignant Properties of Hepatoma Cells*. J. Biol. Chem. 287, 4759–4772 10.1074/jbc.M111.277699

29 Mannowetz, N., Wandernoth, P. and Wennemuth, G. (2012) Basigin interacts with both MCT1 and MCT2 in murine spermatozoa. J. Cell. Physiol. 227, 2154–2162 10.1002/jcp.22949

30 Wanaguru, M., Liu, W., Hahn, B. H., Rayner, J. C. and Wright, G. J. (2013) RH5–Basigin interaction plays a major role in the host tropism of *Plasmodium falciparum*. Proc. Natl. Acad. Sci. 110, 20735–20740 10.1073/pnas.1320771110

31 Arthur, J. S. C., Elce, J. S., Hegadorn, C., Williams, K. and Greer, P. A. (2000) Disruption of the Murine Calpain Small Subunit Gene, *Capn4* : Calpain Is Essential for Embryonic Development but Not for Cell Growth and Division. Mol. Cell. Biol. 20, 4474–4481 10.1128/MCB.20.12.4474-4481.2000

32 Beningo, K. A., Dembo, M., Kaverina, I., Small, J. V. and Wang, Y. (2001) Nascent Focal Adhesions Are Responsible for the Generation of Strong Propulsive Forces in Migrating Fibroblasts. J. Cell Biol. 153, 881–888 10.1083/jcb.153.4.881

33 Dembo, M. and Wang, Y.-L. (1999) Stresses at the Cell-to-Substrate Interface during Locomotion of Fibroblasts. Biophys. J. 76, 2307–2316 10.1016/S0006-3495(99)77386-8

34 Guo, W., Frey, M. T., Burnham, N. A. and Wang, Y. (2006) Substrate Rigidity Regulates the Formation and Maintenance of Tissues. Biophys. J. 90, 2213–2220 10.1529/biophysj.105.070144

35 Tang, Z., Kang, B., Li, C., Chen, T. and Zhang, Z. (2019) GEPIA2: an enhanced web server for large-scale expression profiling and interactive analysis. Nucleic Acids Res. 47, W556– W560 10.1093/nar/gkz430

36 Chang, S. S., Guo, W., Kim, Y. and Wang, Y. (2013) Guidance of Cell Migration by Substrate Dimension. Biophys. J. 104, 313–321 10.1016/j.bpj.2012.12.001

37 Engler, A. J., Sen, S., Sweeney, H. L. and Discher, D. E. (2006) Matrix Elasticity Directs Stem Cell Lineage Specification. Cell 126, 677–689 10.1016/j.cell.2006.06.044

38 Mohammadi, H., Janmey, P. A. and McCulloch, C. A. (2014) Lateral boundary mechanosensing by adherent cells in a collagen gel system. Biomaterials 35, 1138–1149 10.1016/j.biomaterials.2013.10.059

39 Menon, S. and Beningo, K. A. (2011) Cancer Cell Invasion Is Enhanced by Applied Mechanical Stimulation. PLoS ONE (Gullberg, D., ed.) 6, e17277 10.1371/journal.pone.0017277

40 Myers, K. A., Applegate, K. T., Danuser, G., Fischer, R. S. and Waterman, C. M. (2011) Distinct ECM mechanosensing pathways regulate microtubule dynamics to control endothelial cell branching morphogenesis. J. Cell Biol. 192, 321–334 10.1083/jcb.201006009

41 Guilak, F., Cohen, D. M., Estes, B. T., Gimble, J. M., Liedtke, W. and Chen, C. S. (2009) Control of Stem Cell Fate by Physical Interactions with the Extracellular Matrix. Cell Stem Cell 5, 17–26 10.1016/j.stem.2009.06.016

42 Pelham, R. J. and Wang, Y. (1997) Cell locomotion and focal adhesions are regulated by substrate flexibility. Proc. Natl. Acad. Sci. 94, 13661–13665 10.1073/pnas.94.25.13661

43 Plotnikov, S. V., Pasapera, A. M., Sabass, B. and Waterman, C. M. (2012) Force Fluctuations within Focal Adhesions Mediate ECM-Rigidity Sensing to Guide Directed Cell Migration. Cell 151, 1513–1527 10.1016/j.cell.2012.11.034

44 Wolf, K., Te Lindert, M., Krause, M., Alexander, S., Te Riet, J., Willis, A. L., et al. (2013) Physical limits of cell migration: Control by ECM space and nuclear deformation and tuning by proteolysis and traction force. J. Cell Biol. 201, 1069–1084 10.1083/jcb.201210152

45 Shi, J.-H., Huitfeldt, H. S., Suo, Z.-H. and Line, P.-D. (2011) Growth of Hepatocellular Carcinoma in the Regenerating Liver. Liver Transpl. 17, 866–874 10.1002/lt.22325

46 Savarese-Brenner, B., Heugl, M., Rath, B., Schweizer, C., Obermayr, E., Stickler, S., et al. (2022) MUC1 and CD147 Are Promising Markers for the Detection of Circulating Tumor Cells in Small Cell Lung Cancer. Anticancer Res. 42, 429–439 10.21873/anticanres.15501

47 Sun, J. and Hemler, M. E. (2001) Regulation of MMP-1 and MMP-2 Production through CD147/Extracellular Matrix Metalloproteinase Inducer Interactions1. Cancer Res. 61, 2276–2281

48 Zucker, S., Hymowitz, M., Rollo, E. E., Mann, R., Conner, C. E., Cao, J., et al. (2001) Tumorigenic Potential of Extracellular Matrix Metalloproteinase Inducer. Am. J. Pathol. 158, 1921–1928 10.1016/S0002-9440(10)64660-3

49 Yu-li Wang. (2009) Traction forces and rigidity sensing of adherent cells. In 2009 Annual International Conference of the IEEE Engineering in Medicine and Biology Society, pp 3339–3340, IEEE, Minneapolis, MN 10.1109/IEMBS.2009.5333200

50 Ahn, W. S., Jeon, J., Jeong, Y., Lee, S. J. and Yoon, S. K. (2008) Effect of culture temperature on erythropoietin production and glycosylation in a perfusion culture of recombinant CHO cells. Biotechnol. Bioeng. 101, 1234–1244 10.1002/bit.22006

51 Cui, D., Yamamoto, K. and Ikeda, E. (2024) High-Mannose–Type Glycan of Basigin in Endothelial Cells Is Essential for the Opening of the Blood–Brain Barrier Induced by Hypoxia, Cyclophilin A, or Tumor Necrosis Factor-α. Am. J. Pathol. 194, 612–625 10.1016/j.ajpath.2023.11.007

52 Priglinger, C. S., Szober, C. M., Priglinger, S. G., Merl, J., Euler, K. N., Kernt, M., et al. (2013) Galectin-3 Induces Clustering of CD147 and Integrin-β1 Transmembrane Glycoprotein Receptors on the RPE Cell Surface. PLoS ONE (Strauβ, O., ed.) 8, e70011 10.1371/journal.pone.0070011

53. Hunter, G. Basigin associates with integrin in order to regulate perineurial glia and Drosophila nervous system morphology. J Neuro

54 Cui, D., Yamamoto, K. and Ikeda, E. (2024) High-Mannose–Type Glycan of Basigin in Endothelial Cells Is Essential for the Opening of the Blood–Brain Barrier Induced by Hypoxia, Cyclophilin A, or Tumor Necrosis Factor-α. Am. J. Pathol. 194, 612–625 10.1016/j.ajpath.2023.11.007

