## Supporting information for "Regulation of traction force through the direct binding of Basigin (CD147) and Calpain 4"

Supporting Information 1.

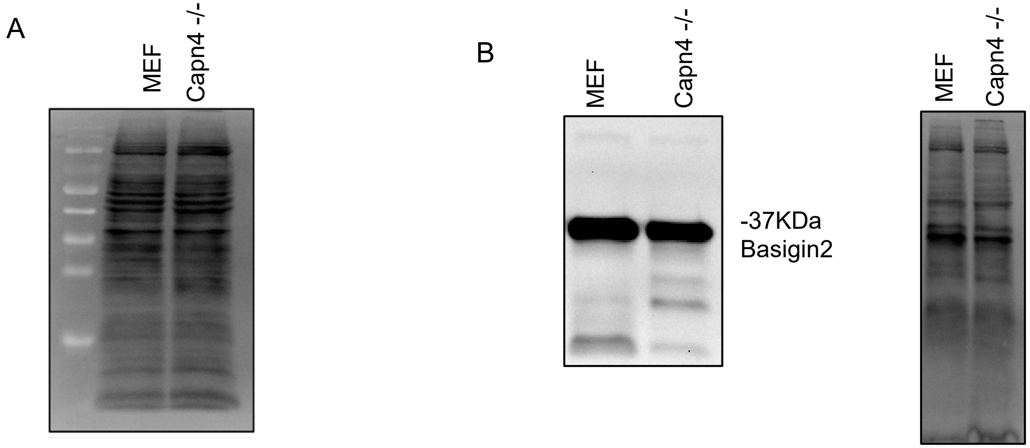


<u>Total Protein and Enzymatic Digestion of Glycosylation sites on Basigin</u>

(A) Total protein for figure 2A.

(B) Western blot probed for basigin 2 and total protein of lysates treated with PNGase. The single strong band represents the low glycosylation form of basigin2.

Supporting Information 2.

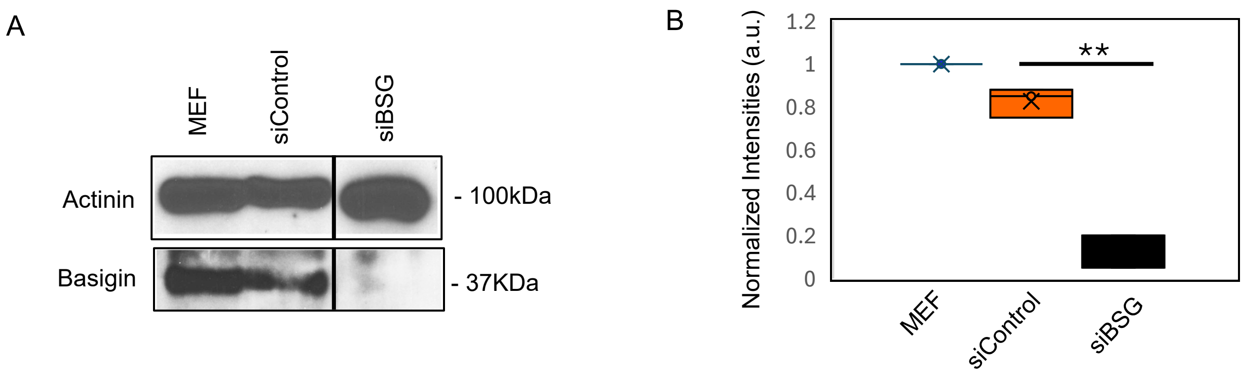


Knockdown of Basigin in Mouse Embryonic Fibroblast

(A) Cell extracts were made 36 hours after nucleofection with siRNA sequence targeting basigin. Western blots probed with anti-basigin antibody had 95% reduction in basigin expression. *B.* Intensities normalized to α-actinin load control. Statistical analysis was performed by student's *t* test. The basigin knockdown lanes were run on the same gel but three lanes apart to allow for maximum volume of the load.

(B) The intensities of the individual lanes were normalized to the load control.
